# Neural components underlying successful free recall are specific to episodic memory

**DOI:** 10.1101/2025.07.25.666835

**Authors:** Riley D. DeHaan, Youssef Ezzyat, Michael J. Randazzo, Aditya M. Rao, Alexander M. Papanastassiou, Aaron S. Geller, Bradley C. Lega, Joshua P. Aronson, Robert E. Gross, Barbara C. Jobst, Kareem A. Zaghloul, Gregory A. Worrell, Sameer A. Sheth, Michael R. Sperling, Michael J. Kahana

## Abstract

Episodic memory depends upon activity distributed across the brain. However, the activity underlying memory has largely been examined within single tasks in isolation. Thus it is unclear to what extent prior findings reflect task-general rather than memory-specific cognitive processes. Here we address this question using data from 371 patients recorded intracranially who performed a free recall task with encoding and retrieval phases alongside an arithmetic distractor phase. We ask whether neural decoders fit to predict behavior from one phase transfer to the others. Encoding-retrieval transfer exceeds both arithmetic-encoding and arithmetic-retrieval transfer and therefore cannot be explained solely by processes supporting arithmetic. We further detect transfer between arithmetic and retrieval but not between arithmetic and encoding. The brain-behavioral relations observed in these tasks thus do not merely reflect a single task-general factor of activity. We propose cross-task decoding as a method for identifying the neural factor structure underlying distinct cognitive processes.

## Introduction

Cognitive neuroscience aims to relate brain activity to cognitive processes. By predicting outcomes during a behavioral task putatively invoking a cognitive process, neural decoders provide a powerful tool for establishing brain-behavior relations. However, neuroscientists frequently use decoders to contrast conditions within a single cognitive task, leaving open the possibility that the observed brain-behavior relations correspond to task-general rather than task-specific processes. Activity recorded during any individual task will be a mixture of activity related to the process under study and other processes not of immediate interest, challenging interpretations of neural effects.

This problem arises frequently within the field of human memory, in which univariate and multivariate methods have established robust relations between widespread activity measured during item study and subsequent recall success (Wagner et al., 1998; Ezzyat et al., 2017). Whereas these subsequent memory effects have been observed in brain regions specific to memory function such as the hippocampus, they have also been detected in sensory areas involved in vision and frontal regions implicated in executive function (Kim, 2011; Long et al., 2014; Urgolites et al., 2020). However, by comparing subsequently remembered and non-remembered items in the context of a single memory task, the majority of prior studies cannot distinguish task-specific contributions to memory encoding from task-general processes such as attention or arousal.

Recent work has investigated the neural relation between encoding and the related but distinct process of memory retrieval, following long-standing questions in the field (Tulving, 1979). In line with the idea that the same neural substrates engaged with processing memory contents at the time of encoding also play a role in storing those memories, item-specific activity patterns present at the time of encoding have been found to be reinstated at the time of retrieval (Polyn et al., 2005; Manning et al., 2011; Gordon et al., 2014; Tompary et al., 2016). In addition to item-specific patterns of neural similarity between encoding and retrieval, subsequent memory effects have also been related to activity predicting successful retrieval in comparison to periods of failed memory search when averaged across items. Kragel et al. (2017) in particular find that multivariate neural decoders fit to predict subsequent recall from activity during encoding also distinguish brain activity during successful retrieval from activity at moments of failed memory search during the recall phase. Similarly, neural decoders fit to predict retrieval success from activity during the recall phase generalize to predict success during encoding. Whereas these results suggest shared neural processes underlying successful encoding and retrieval, they are unable to determine whether the shared activity is specific to memory processing or instead reflects more general cognitive functions engaged during not only encoding and retrieval but also tasks unrelated to memory.

Here we explore the question of isolating task-specific and task-general components of neural activity predicting performance in human memory. Using a large historical dataset of the free recall experiment collected in 371 intracranially recorded epileptic patients, we compare neural activity predicting behavioral outcomes during encoding and retrieval to that of an arithmetic distractor task placed between the encoding and retrieval phases. By contrasting encoding and retrieval against a third cognitive task, we are able to distinguish whether the relation between encoding and retrieval is characterized by a single task-general factor encompassing all three tasks, or whether multiple neural factors are required to account for relations between neural activity across these tasks. If indeed neural activity associated with behavior during these three tasks cannot be described using a single factor model, this approach further allows for addressing the questions of whether arithmetic-related activity overlaps with activity during encoding or separately with activity during retrieval. Comparing across multiple cognitive tasks may thereby further our understanding of the relation between encoding and retrieval processes.

Using an arithmetic distractor as our comparison task offers several benefits. Arithmetic problem solving is known to engage task-general cognitive processes including working memory, executive function, and sustained attention. Arithmetic generates activity in prefrontal cortex as well as superior and inferior parietal lobules (Arsalidou and Taylor, 2011), and math performance has been associated with performance on tests of executive functioning and with lesions to frontal areas involved in executive function and attention (Cragg and Gilmore, 2014; Lucchelli and De Renzi, 1993). Furthermore, we expected the distractor would largely not engage encoding processes. Subjects are not instructed to remember distractor problems. Indeed, considering distractors have large effects on recall rates by abolishing the recency effect present in immediate free recall, subjects are disincentivized from devoting cognitive effort to remembering these problems (Kahana, 2012). These properties make arithmetic useful for differentiating task-specific and task-general neural factors predicting memory behavior. Additionally, our math task’s placement as a distractor embedded within a free recall experiment ensured subjects provided data during both mnemonic and arithmetic-related processing within the same sessions. This design minimized reductions in neural similarity between tasks due to drift in recording properties as may have occurred had subjects completed these tasks in separate sessions.

Our three-way cross-task approach allows us to answer the question of whether activity predicting successful memory behavior is specific to memory function or else better explained by task-general processes also involved in arithmetic. We first demonstrate that intracranial recordings reliably decode math responses. Univariate anatomical analyses additionally reveal specific effects within frontal regions during the math task in line with a central role for executive function. We then ask whether decoders fit to predict behavior within a given task phase generalize to predicting outcomes in the other phases as a measure of task-related neural similarity. Our results indicate shared neural factors between encoding and retrieval and between retrieval and arithmetic. However, we find no evidence for a factor shared between encoding and math, suggesting that a single task-general factor of activity cannot account for the observed relation between these tasks. We further uncover significantly greater neural similarity between encoding and retrieval than between either memory task and math, underscoring our claim that activity shared between encoding and retrieval cannot be attributed solely to processes engaged during arithmetic. Together these findings suggest that the activity predicting mnemonic behavior reflects memory-specific processes and illustrate the benefits of comparing neural activity across multiple distinct tasks within the same subjects.

## Results

We asked whether shared patterns of neural activity that predict successful memory during both encoding and retrieval could be accounted for by a single task-general factor of activity that would also predict behavior during cognitive tasks unrelated to memory. Alternatively, activity underlying encoding and retrieval may be specific to memory and therefore not generalize to other tasks. To address this question, we used multivariate classification to decode variability in mnemonic and cognitive task performance. Subjects studied lists of 12 common words, performed a mental arithmetic task, and then attempted free recall of the studied items (see Figure 1A). We first assessed whether spectral EEG features recorded from widespread brain regions can reliably decode behavior during each task phase: Encoding, Retrieval, and Math. Specifically, for Encoding we contrast activity between study periods for remembered and not remembered items from item presentation onset to 1366 ms after onset. For Retrieval, we compare the interval from 600 to 100 ms before the retrieval vocalization to matched silent periods, or “deliberations”, during the recall phase indicative of failed memory search as in prior work (Burke et al., 2014; Kragel et al., 2017; see Methods). For Math, we contrast activity in the moments before typed arithmetic problem responses between slower and faster response times (based on a median split) as a measure of arithmetic task engagement. While we considered contrasting correct and incorrect math responses, subject accuracy was too high to ensure sufficient incorrect responses for stable decoder training, with a median accuracy across exploration subjects of 95.4%. To reduce the influence of motor-related activity on any neural similarity observed between retrieval and math, we used spectral epochs during Math from 1800 to 500 ms before subject responses, distancing our analysis epochs from task-related motor movements. We also excluded electrodes in cortical areas related to somatosensation, motor function, and speech from our analyses (see Methods). For each epoch and electrode, Morlet wavelets provided 8 spectral iEEG powers logarithmically spaced from 5 Hz to 175 Hz. These wavelet powers were averaged across time to generate the frequencies-by-electrodes spectral features for a given epoch. After presenting neural decoding within each task phase, we report the univariate features that differentiate task responses across regions of interest implicated in memory and mathematical cognition (see Figure 1B for electrode coverage). Having analyzed the neural features particular to each task, we examine whether models trained on one task phase generalize to predict performance during other task phases in order to elucidate the shared and distinct neural factors relating these tasks.

**Figure 1:**
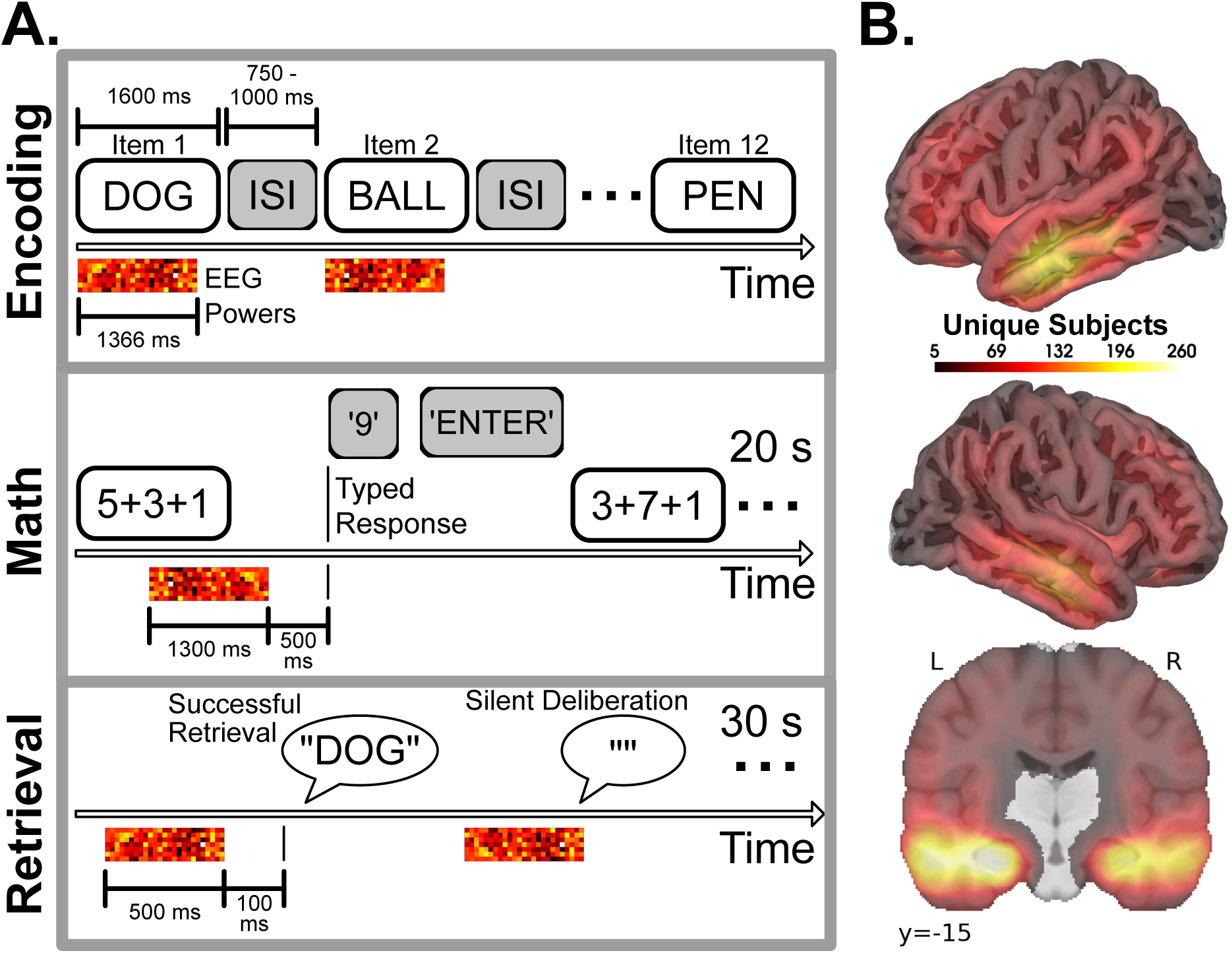
Brain recordings during mnemonic and math tasks. A. Schematic of the free recall experiment. Subjects studied lists of 12 common words (Encoding) which they subsequently recalled after a delay (Retrieval). During the delay period, subjects performed a mental arithmetic task (Math). Spectral iEEG powers from 5 Hz to 175 Hz at each electrode were computed across event epochs defined for each task (colored matrices). B. Electrodes recorded brain activity while subjects performed the memory and math tasks. Brain heat maps illustrate counts of subjects with electrode coverage within 5 mm of a voxel across all exploration (*N* = 195) and confirmation (*N* = 176) subjects (not all subjects contributed data for all task phases; see Methods for exclusion criteria).

We developed our primary analyses of within- and cross-task neural decoding and neural similarity in roughly half of our data. We subsequently preregistered these analyses before replicating them on the held-out half (see Methods). We report results separately for our exploration and confirmation datasets. Not all analyses were preregistered, and we explicitly identify exploratory analyses throughout our results.

### Within-task neural correlates of encoding, retrieval, and arithmetic

We first examined the multivariate and univariate neural correlates of behavioral outcomes specific to the three task phases: Encoding, Retrieval, and Math. Prior work has shown robust effects of mnemonic success on widespread neural activity during both encoding and retrieval (Ezzyat et al., 2017; Kragel et al., 2017; Weidemann et al., 2019). We extend these results by demonstrating reliable classification of trials with slower compared to faster arithmetic problem responses as measured by Area Under the Curve (AUC) using ridge-penalized logistic regression classifiers. As shown in Figure 2A, we observe significant within-task decoder generalization to held-out sessions for each of the three tasks (one-sample t-tests against random chance of AUC = 0.5: confirmation: encoding *t*_60_ = 14.1, *p* < 10^−19^; retrieval, *t*_60_ = 18.5, *p* < 10^−25^; math *t*_60_ = 9.8, *p* < 10^−13^; exploration: encoding *t*_73_ = 12.8, *p* < 10^−19^; retrieval, *t*_73_ = 17.6, *p* < 10^−27^; math *t*_73_ = 12.8, *p* < 10^−19^). Neural activity further demonstrates reliable effects of behavioral outcomes during these task phases when resolved across spectral frequencies and major regions of interest related to memory and mathematical cognition (Figure 2B, C, D; these univariate analyses were not preregistered but are presented separately for the exploration and confirmation data for comparison). Whereas increased high-frequency activity and decreased low-frequency activity, or *spectral tilt*, across widespread brain areas mark successful encoding and imminent correct retrieval, a seemingly similar pattern of spectral features predicts slower correct response times on our arithmetic task. Although the spectral tilt pattern appeared for all three contrasts, it emerged in somewhat different regions across task. These spectral tilt effects during slower math problems appear in frontal regions associated with tasks involving executive function. These findings align with work showing selective frontal activity for arithmetic problem solving requiring procedural rather than strictly retrieval-based strategies given the additional complexity of our math task in having subjects sum three operands compared to two in more common arithmetic task designs (Sokolowski et al., 2023; Ashcraft, 1992). In addition, the observation that slower rather than faster problems exhibit increased frontal high-frequency activity may be explained by differences in math problem difficulty. More difficult problems may lead to slower arithmetic performance and greater spectral tilt suggestive of greater cognitive demands. Alternatively, this spectral tilt effect may be interpreted in terms of an endogenous state of cognitive efficiency. On some problems, subjects happen to retrieve or calculate an answer quickly and with minimal neural processing, leading to the observed effect. These possibilities motivated with a control analysis accounting for problem difficulty we present below (see section *Controlling for arithmetic problem difficulty*). In line with these interpretations, we coded our neural contrasts of arithmetic problem as slower responses minus faster responses to provide a measure of arithmetic task engagement.

**Figure 2:**
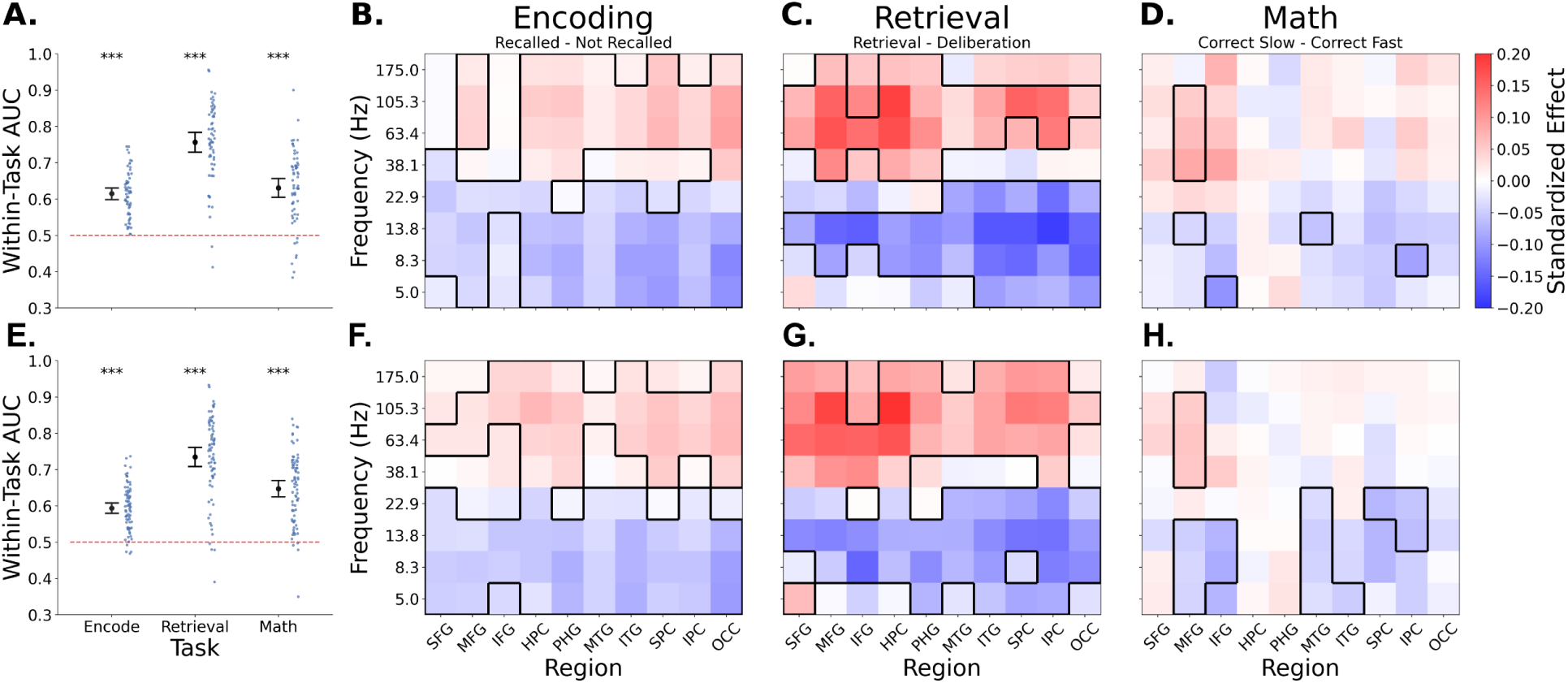
Intracranial electrophysiology predicts successful episodic memory and math engagement. Top row presents results from our confirmation sample: A. Classification AUC for decoding cognitive outcomes during the Encoding, Retrieval, and Math phases of the free recall experiment. Dots show subject classifier scores. B., C., D. Univariate behavioral contrasts of neural spectral power in Encoding (B.), Retrieval (C.), and Math (D.) across regions of interest and spectral frequency. These univariate analyses were not preregistered. Colors indicate Hedge’s *g* with borders indicating significant ROI-frequency combinations (p < 0.05 FDR-corrected across frequencies and ROIs). Bottom row (E., F., G., H.) presents the same results in our exploration sample. AUC: Area Under the Curve; STG, MFG, IFG: superior, middle, and inferior frontal gyri; PHG: parahippocampal gyrus; HPC: hippocampus; MTG, ITG: middle and inferior temporal gyri; IPC, SPC: inferior and superior parietal cortex; OCC: occipital cortex. Error bars show standard 95% confidence intervals around the mean. ***: *p* < 0.001, **: *p* < 0.01, *: *p* < 0.05, n.s.: *p* ≥ 0.05.

### Cross-task decoding reveals memory-specific activity

We asked whether neural decoders fit to predict arithmetic problem-solving responses would transfer to predicting success during both encoding and retrieval of episodic memories and vice versa. If so, we hypothesized that neural activity shared between encoding and retrieval could reflect a single neural factor common to all three tasks. On the other hand, Math may generalize to Encoding or Retrieval but not both, suggesting a dissociation in the neural activity engaged during Encoding and Retrieval. Figure 3 illustrates cross-task decoding among Encoding, Retrieval, and Math classifiers. We expected, based on prior work (Kragel et al., 2017), to find reliable transfer between episodic memory Encoding and Retrieval. Indeed, decoders trained on Retrieval reliably predicted Encoding success (one-sample t-test against AUC=0.5: confirmation: *t*_60_ = 12.2, *p* < 10^−17^; exploration: *t*_73_ = 13, *p* < 10^−19^) and vice versa (confirmation: *t*_60_ = 14.2, *p* < 10^−20^; exploration: *t*_73_ = 13.4, *p* < 10^−20^). Of greater interest, however, was the finding that Math task decoders reliably predicted behavioral outcomes during Retrieval trials (confirmation: *t*_60_ = 2.9, *p*_FDR_ = 0.025; exploration: *t*_73_ = 5.1, *p*_FDR_ < 10^−4^) and vice versa (confirmation: *t*_60_ = 3.37, *p*_FDR_ = 0.011; exploration: *t*_73_ = 4.9, *p*_FDR_ < 10^−4^). In contrast, we did not observe transfer from Math to Encoding (confirmation: *t*_60_ = −0.42, *p*_FDR_ = 1.0; exploration: *t*_73_ = 0.51, *p*_FDR_ = 1.0) or from Encoding to Math (confirmation: *t*_60_ = 0.12, *p*_FDR_ = 1.0; exploration: *t*_73_ = 0.82, *p*_FDR_ = 1.0). Critically we find significantly greater transfer from math to retrieval than from math to encoding (paired t-test: confirmation: *t*_60_ = 3.7, *p*_FDR_ = 0.0012; exploration: *t*_73_ = 5.3, *p*_FDR_ < 10^−5^) and greater transfer from retrieval to math than from encoding to math (confirmation: *t*_60_ = 3.69, *p*_FDR_ = 0.0012; exploration: *t*_73_ = 4.1, *p*_FDR_ = 0.00032). To account for the effect of within-task decoding on these differences in cross-task decoding, we extended these comparisons after normalizing the cross-task scores by the within-task score for the test task (see Methods; this analysis was not preregistered). These normalized scores reflect the proportion of signal that transfers relative to the within-task effect size. After controlling for within-task decoding strength, we continue to observe significantly greater transfer from math to retrieval than from math to encoding (paired t-test: confirmation: *t*_40_ = 2.93, *p*_FDR_ = 0.010; exploration: *t*_42_ = 2.66, *p*_FDR_ = 0.020) and greater transfer from encoding to retrieval than from encoding to math (confirmation: *t*_40_ = 8.43, *p*_FDR_ < 10^−8^; exploration: *t*_42_ = 6.92, *p*_FDR_ =< 10^−6^; see Supplementary Figure 1).

**Figure 3:**
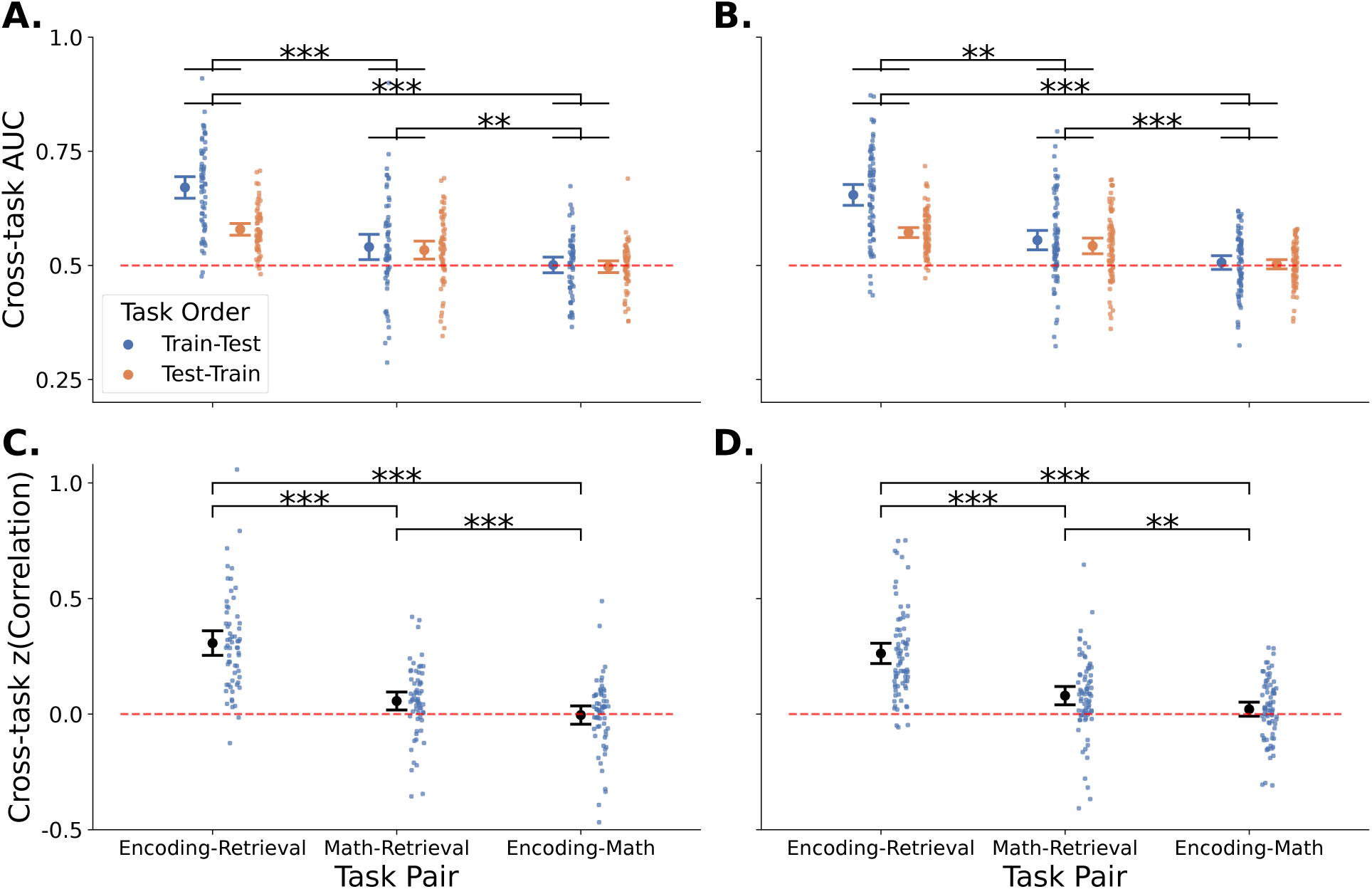
Cross-task decoding isolates neural factors underlying encoding, retrieval, and math. Here we evaluate how neural decoders trained on a given task predict performance in another task. A., B. Cross-task decoding between item encoding, spontaneous retrieval, and math distractor performance during free recall (A: confirmation data, B: exploration data). Significance bars compare pairs of tasks, with significance indicators based on the maximum p-value obtained across the two directions of cross-task decoding for FDR-corrected paired t-tests (see main text). C., D. Pearson correlations between within-task contrasts of cognitive success (C: confirmation data, D: exploration data). Dots show subject-level statistics. Error bars display standard 95% confidence intervals around the mean. Significance bars reflect FDR-corrected paired t-tests. ***: *p* < 0.001, **: *p* < 0.01, *: *p* < 0.05, n.s.: *p* ≥ 0.05.

This selective pattern of cross-task neural decoding implicates distinct components of shared variability between encoding and retrieval and between math and retrieval that are not shared between encoding and math. We further find greater transfer from encoding to retrieval than from math to retrieval (paired t-test: confirmation: *t*_60_ = 7.9, *p*_FDR_ < 10^−9^; exploration: *t*_73_ = 7.3, *p*_FDR_ < 10^−8^) and greater transfer from retrieval to encoding than from retrieval to math (confirmation: *t*_60_ = 4.0, *p*_FDR_ = 0.00080; exploration: *t*_73_ = 3.0, *p*_FDR_ = 0.0083). These observations indicate a stronger correlation between encoding and retrieval than between these memory tasks and math, suggesting the activity shared between encoding and retrieval is not explained by processes engaged during arithmetic and instead is specific to episodic memory.

To assess whether these results stem from our particular choice of neural similarity measure, we performed an additional comparison using a form of representational similarity analysis (Kriegeskorte et al., 2008). Specifically, we computed correlations across tasks between subject-level vectors of univariate standardized task effects. These task effects were quantified by Hedge’s *g* for each contact and frequency. This approach yielded an identical pattern of results to the cross-task neural decoding analysis above (Figure 3C, D). The mean correlation across subjects between neural effect vectors for encoding and retrieval (confirmation: *t*_60_ = 11.3, *p* < 10^−15^; exploration: *t*_73_ = 11.7, *p* < 10^−17^) and for retrieval and math significantly exceeded random chance (confirmation: *t*_60_ = 2.8, *p*_FDR_ = 0.019; exploration: *t*_73_ = 4.0, *p*_FDR_ = 0.0005). However, this relation was not found between encoding and math (confirmation: *t*_60_ = −0.2, *p*_FDR_ = 1.0; exploration: *t*_73_ = 1.4, *p*_FDR_ = 1.0). We find the mean correlation to be significantly greater both for encoding and retrieval (confirmation: *t*_60_ = 9.8, *p*_FDR_ < 10^−9^; exploration: *t*_73_ = 10.6, *p*_FDR_ < 10^−14^) and for retrieval and math compared to encoding and math (confirmation: *t*_60_ = 3.8, *p*_FDR_ = 0.00061; exploration: *t*_73_ = 3.4, *p*_FDR_ = 0.002). We also find the correlation between encoding and retrieval exceeds the correlation between retrieval and math (confirmation: *t*_60_ = 7.8, *p*_FDR_ < 10^−9^; exploration: *t*_73_ = 6.7, *p*_FDR_ < 10^−7^). These results indicate that our cross-task decoding results are not an artifact of our particular measure of neural similarity and support our finding of two distinct neural factors, one shared between encoding and retrieval and another between retrieval and math.

### Controlling for arithmetic problem di**ffi**culty

Our finding that spectral tilt predicted slower rather than faster arithmetic responses suggested two interpretations. Response times and spectral tilt are both driven by 1. problem difficulty or by 2. endogenous states of neural activity leading to some problems being solved quickly and with minimal cognitive effort. To determine whether problem difficulty can account for our results, we repeated our previous analyses after controlling for variability in math problem difficulty using a mixed-effects model of math problem RTs fit to the behavioral data across all subjects fit separately for our exploration and confirmation data. Our model of problem difficulty included several features of summation problems, including problem sum and whether the problem required carry operation(s) (see Methods for all model factors). This model explained 21.3% of the residual variance in logarithmic RTs after accounting for subject and session intercepts in our confirmation data (exploration: 22.4%). Figure 4A presents average problem RT as a function of problem accuracy across all problem presentations while C and D present average RTs and model predictions for each problem respectively.

**Figure 4:**
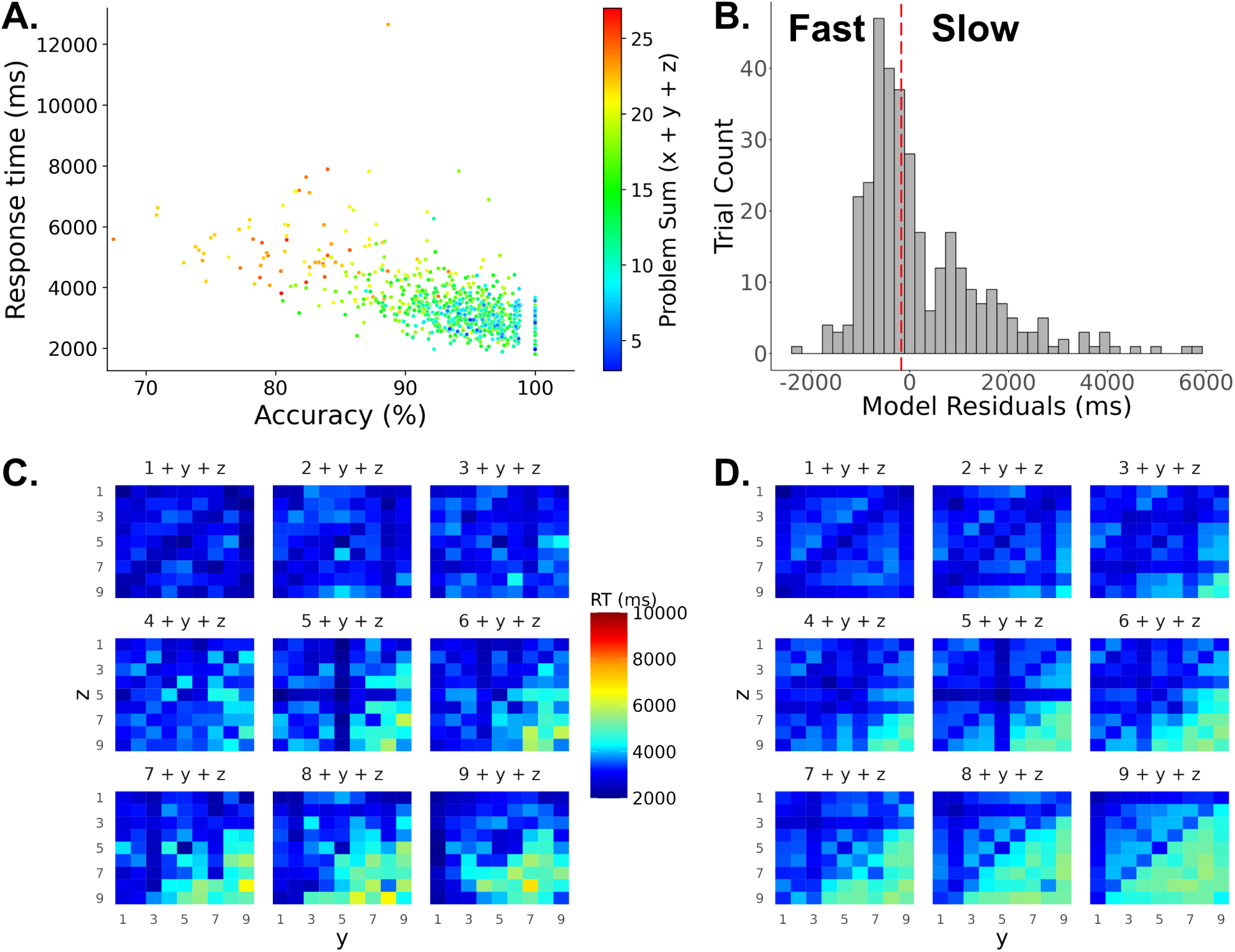
**Controlling for math problem di**ffi**culty.** A. Response time vs. accuracy for each of the presented 729 math problems averaged across trials. B. Approach to controlling for problem difficulty in our cross-task decoding comparisons. A trial was labeled “slow” if its prediction residual was slower than the median residual for that session and “fast” otherwise. C., D. Response times for each problem averaged across trials (C.) and model predicted response times (D.) in our confirmation data.

To control for problem difficulty in our neural analyses, we computed RT residuals from the behavioral model and labeled each math trial as “slow” if its residual was above the residual median for the session and “fast” otherwise (Figure 4B). We then repeated our prior analyses of within-task and cross-task neural decoding. Within-task neural decoders reliably predict adjusted arithmetic performance across subjects (Figure 5A; confirmation: *t*_60_ = 11.1, *p* < 10^−16^; exploration: *t*_73_ = 10.7, *p* < 10^−15^). The univariate neural effects still show increased spectral tilt for slower responses after difficulty adjustment (Figures 5B, C). Critically, cross-task decoding continued to hold between encoding and retrieval (all *p*’s < 10^−17^), from math to retrieval (confirmation: *t*_60_ = 3.3, *p* = 0.014; exploration: *t*_73_ = 4.3, *p*_FDR_ = 0.0002), and from retrieval to math (confirmation: *t*_60_ = 2.9, *p*_FDR_ = 0.023; exploration: *t*_73_ = 5.1, *p*_FDR_ < 10^−4^) as shown in Figure 6. However as before, we find no evidence for cross-task decoding between encoding and math (all *p*’s > 0.6). The encoding-retrieval and retrieval-math cross-task decoding remains significantly greater than the encoding-math cross-task decoding (all *p*’s < 0.002). We obtained analogous results with correlation similarities. These results suggest our findings cannot be entirely attributed to arithmetic problem difficulty. Instead, we suggest an endogenous cognitive state, independent of problem difficulty, also contributes to variability in math responses explained by neural activity.

**Figure 5:**
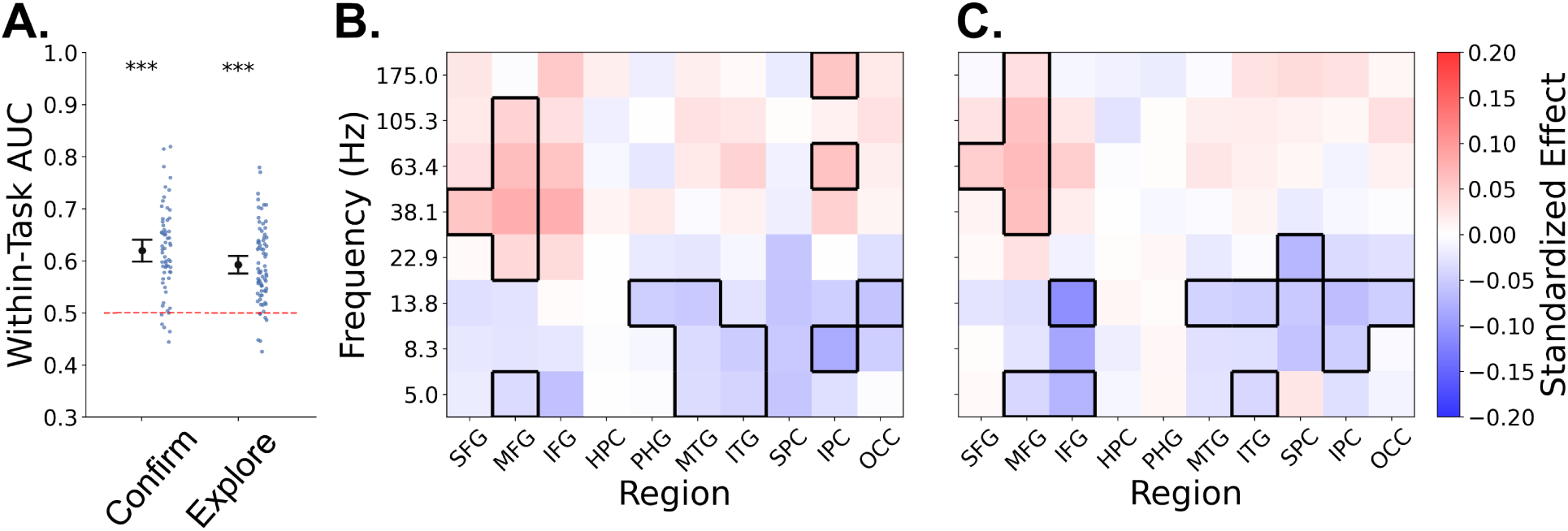
**Electrophysiology still predicts math engagement after controlling for arithmetic problem di**ffi**culty.** A. Neural decoding of Math outcomes after adjusting for problem difficulty shown separately for the confirmation and exploration data. B., C. Univariate features after difficulty adjustment in the confirmation (B.) and exploration (C.) data. Format otherwise follows Figure 2.

### Testing for e**ff**ects of rehearsal on neural similarity between math and retrieval

A potential confound of our findings is subjects rehearsing encoded items during the math distractor. Subject rehearsal could be expected to lead to higher recall rates, retrieval-related activity during the math distractor, and slower math RTs by interfering with problem solving. If so, one could observe spuriously similar activity during the math and retrieval phases unrelated to mathematical cognition. However in this case, we would expect slower math RTs on lists with higher recall rates. We find in contrast that mean list-level math RTs and recall rates do not correlate positively across lists at the population level (Figure 7A; t-test of z-transformed subject correlations against zero: confirmation: *t*_60_ = −3.0, *p* = 0.0035, two-sided objective Bayes factor: *BF*_10_ = 10.2; exploration: *t*_70_ = −1.6, *p* = 0.11, *BF*_10_ = 0.60). We include Bayes factors in these analyses to allow for showing evidence for the null hypothesis of no rehearsal confound. These subject-level correlations additionally do not themselves correlate with subject-level cross-task decoding scores between retrieval and math in either direction of task transfer (confirmation: retrieval-to-math: *r* = 0.026; math-to-retrieval: *r* = 0.0021; exploration: retrieval-to-math: *r* = 0.085; math-to- retrieval: *r* = 0.156; all p’s > 0.1 and all Bayes factors < 0.25). We find analogous results comparing list-level recall rates to mean math RT residuals from our model of problem difficulty (Figure 7B). Rehearsal during the distractor thus cannot account for the observed pattern of cross-task decoding.

**Figure 6:**
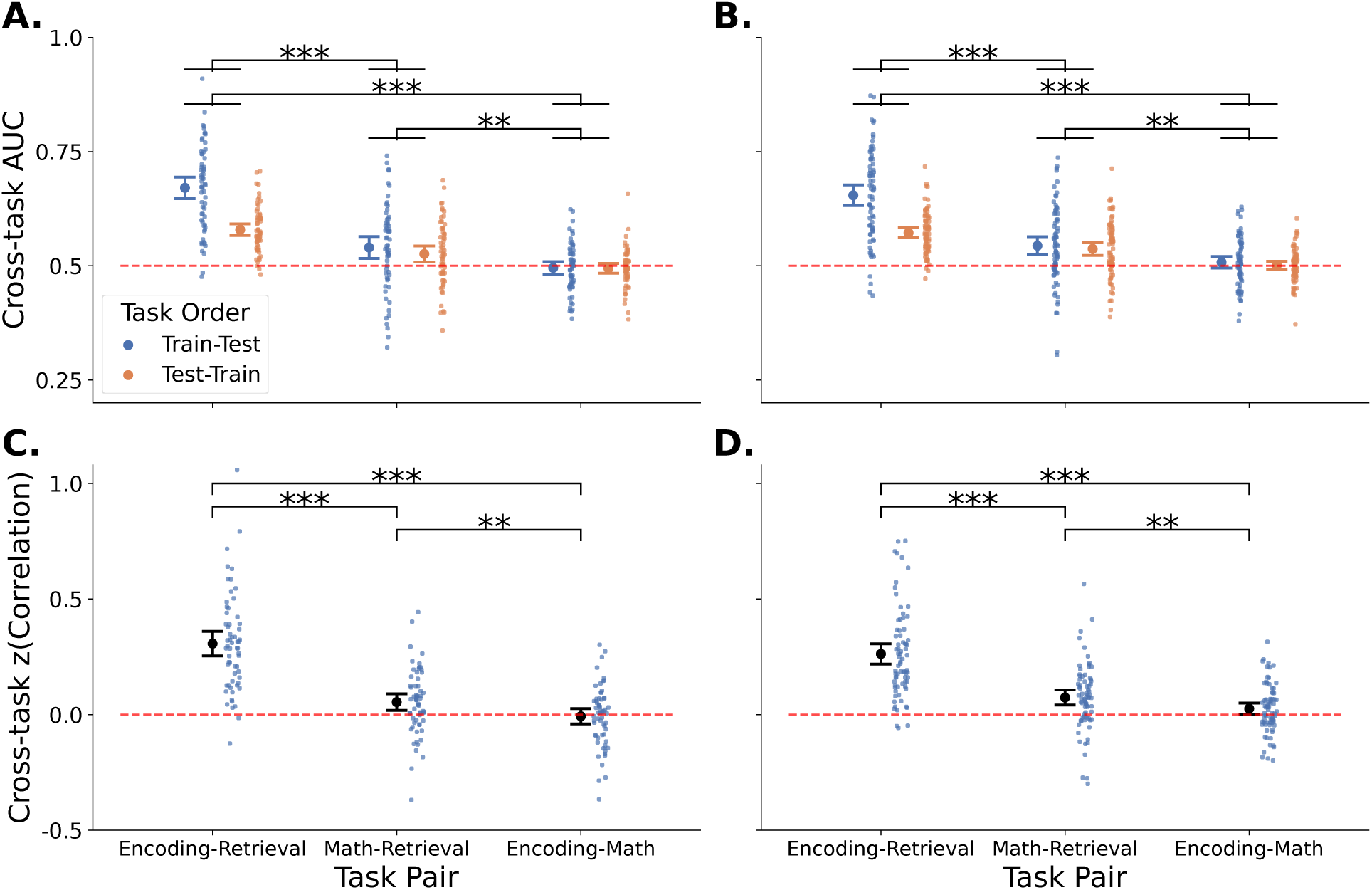
**Cross-task neural factor structure holds after adjusting for problem di**ffi**culty.** After adjusting the math response labels for problem difficulty, we reevaluate how classifiers trained on a given task predict performance in another task. Format follows Figure 3.

**Figure 7:**
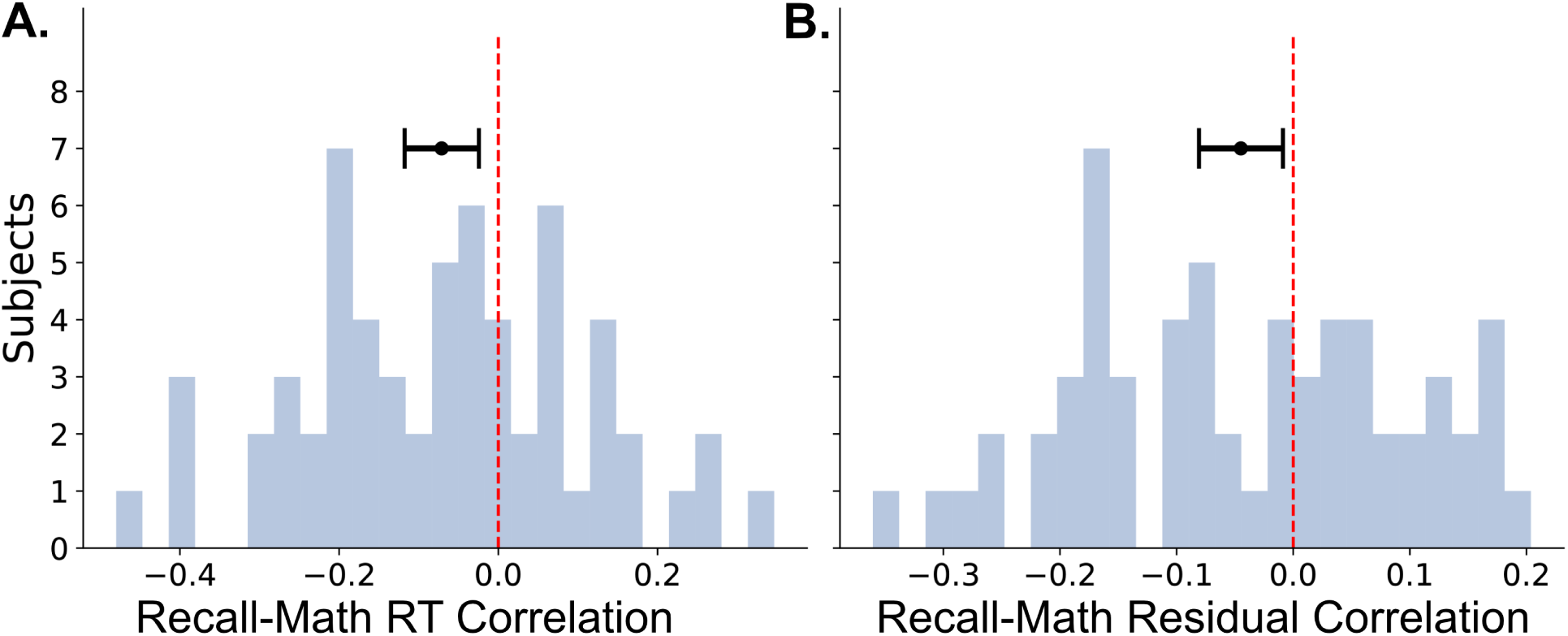
**Memory rehearsal cannot explain neural similarity between retrieval and math**. A. Subject-level correlations between list-level recall rates and average math RTs for the first three math responses in each list. Results show subjects included in the cross-task decoding analyses for the confirmation data. B. Same as A., but using math RT residuals from the statistical model of problem difficulty in place of RTs. Error bars show 95% standard confidence intervals around the mean.

## Discussion

We asked whether neural activity predicting successful memory encoding and retrieval can be primarily attributed to a factor of activity specific to memory processing or to a task-general factor also involved in behavior unrelated to memory. To address this question, we compared the encoding and retrieval periods during a free recall experiment with an arithmetic problem solving task known to require attentional and executive control processes. Participants performed this math task as a distractor between the encoding and retrieval periods. Using a dataset comprising intracranial recordings from 371 patients, we first demonstrated that math response times could be reliably decoded from activity just prior to problem response. We then asked whether neural decoders trained on each task generalize to the other tasks. These cross-task decoding analyses allowed us to evaluate whether a single factor or multiple factors underlie neural decoding within these tasks. After replicating prior work showing that activity predicting successful encoding also predicts success during retrieval and vice versa, we find that math decoders transfer to retrieval and that retrieval decoders transfer to math. However, math classifiers failed to transfer to encoding and encoding classifiers failed to transfer to math. These results hold after adjusting for within-task decoding performance as well as math problem difficulty and cannot be explained by rehearsal processes occurring during the math distractor task.

Our findings argue against a single-factor, task-general account of neural activity predicting behavioral outcomes in these three cognitive tasks. Rather, our results appear consistent with a two factor model, with one factor shared between encoding and retrieval, and a second factor shared between retrieval and math. A parsimonious interpretation would suggest that the shared features underlying cross-task decoding between encoding and retrieval reflect processes largely specific to memory function. We found significantly stronger cross-task decoding between encoding and retrieval than between either of those tasks and the math distractor. This result indicates the relation between encoding and retrieval cannot be considered merely a byproduct of task-general processes engaged during math. These findings extend the work of Kragel et al. (2017), who previously reported shared activity predicting success between encoding and retrieval in a subset of the data we present. However, they were unable to determine whether this shared component was driven by a general attentional process that would lead to transfer across a wide range of behavioral tasks or by a process specific to memory. By comparing the encoding and retrieval tasks to a math task, we address this limitation.

In addition to the prominent shared activity between encoding and retrieval, we identified a secondary–but reliable–component of neural similarity between retrieval and the math task. We tentatively interpret this second factor as reflecting task-general executive control processes, likely mediated by frontal cortical regions. Supporting this view, slower math responses during putatively more difficult problems showed greater high-frequency activity and decreased low-frequency activity in middle frontal gyrus. Although frontal activation was also present during encoding, the more spatially concentrated effects qualitatively observed during retrieval and math suggest a distinct frontal control-related pattern underlying the component shared between retrieval and math. These findings are consistent with prior work linking both mathematical cognition (Knops et al., 2009; Sokolowski et al., 2023; Pinheiro-Chagas et al., 2024) and episodic retrieval (Cabeza et al., 2003; Badre and Wagner, 2007; Marklund et al., 2007; Barredo et al., 2015; Vatansever et al., 2021) to executive function in frontal regions, particularly for effortful problem or retrieval responses.

In contrast, encoding–retrieval similarity reflected a broader spectral tilt effect encompassing medial and lateral temporal, parietal, and frontal regions. While spectral tilt was also observed during math, particularly in middle frontal gyrus, it was less spatially consistent than in encoding and retrieval. Notably, despite prior findings implicating superior and inferior parietal lobules in mathematical cognition (Grabner et al., 2009; Harvey et al., 2013; Pinheiro-Chagas et al., 2024), our univariate analyses did not reveal consistent parietal effects during the math task. This discrepancy may reflect the strong spatial heterogeneity in effects previously found within parietal and other regions during mathematical processing (Daitch et al., 2016; Pinheiro-Chagas et al., 2024). Despite the less consistent univariate effects for math within individual ROIs, our multivariate decoding approach revealed comparable within-task effect sizes for encoding and math. This observation highlights a benefit of multivariate decoding approaches in capturing local variability in effects commonly observed across neighboring intracranial recording contacts–local variability which would be obscured by averaging across electrodes within an ROI (Jacobs and Kahana, 2009; Daitch and Parvizi, 2018). Ultimately, our multivariate approach was critical in distinguishing the neural factor shared between encoding and retrieval from math-related activity. This approach also allowed us to identify a second shared factor comprising math and retrieval that we argue may reflect task-general control processes.

An alternative interpretation is that the similarity between math and retrieval is less a function of task-general executive control and attentional processes being present during retrieval, but rather is indicative of retrieval processes occurring during math. The addition problems of the form “A + B + C” used in the math distractor phase can be solved by chaining together retrievals of rotely memorized arithmetic facts. For instance, when adding 3 + 2 + 9, a subject could first retrieve the memorized result of the subproblem 3 + 2 = 5 before adding 5 + 9 with the carry-over operation. While solving these arithmetic problems may involve retrieval, it would seem less likely to involve effortful encoding given subjects were not instructed to remember the problems. Retrieval intrinsically embedded within the math task could partially explain any observed transfer between math and retrieval. However, it appears unlikely these retrievals would explain the entirety of the observed transfer effect, given the failure to see similar transfer to the encoding phase when the retrieval phase itself exhibited robust transfer to encoding. Additionally, the univariate regional activations observed during the math phase do not include canonical memory-related regions such as the hippocampus that activated robustly during retrieval and encoding. These observations suggest that the factor shared between retrieval and math is best attributed to task-general processes rather than processes specific to memory, though retrieval of arithmetic facts may also contribute.

As discussed above, our ability to make strong inferences about cognitive processes engaged by the identified neural factors is limited. Experimental tasks generally involve multiple processes to varying degrees, and cross-task comparisons will reflect the combinations of processes present. Ideal comparison tasks will include as few processes as possible. In particular, while we took several precautions to exclude motor- and somatosensory-related activity from our analyses, we cannot fully rule out the possibility that our results are partly driven by motor-planning responses present during both retrieval and math. Additionally, it is possible that the smaller within-task effect sizes of decoding observed in encoding and math compared to retrieval may have limited our precision to observe neural similarity between these task phases. Our findings suggest that a weaker factor shared between encoding and math is unlikely to explain the shared features between encoding and retrieval considering the substantially stronger similarity between these memory tasks. Furthermore we observe the cross-task decoding between encoding and retrieval and between retrieval and math exceeds that between encoding and math after controlling for within-task decoding strength. Although we cannot entirely dismiss the possibility of a factor shared between encoding and math, we find that two predominant factors shared between encoding and retrieval and between retrieval and math best explain our results.

Future studies may address these concerns using optimized distractor tasks. Our distractor included three operands in contrast to more standard arithmetic tasks involving only two (Ashcraft, 1992; Pinheiro-Chagas et al., 2024). Extending our study using a distractor with two operands would shrink the space of relevant problem solving strategies, presumably reducing variability across subjects and problem types and thereby improving precision. In addition, an optimized task design could require subjects to initially press the same key on each trial to indicate they have solved the problem before typing their numerical response, thereby minimizing differences in typing movements across problem responses. Alternatively, subjects could vocalize responses during the math distractor task to be better matched to the retrieval phase. However, the fact that we observe neural similarity between math and retrieval despite them having different motor responses and stimulus sets suggests robustness of the effect and a cognitive explanation rather than one based on stimulus or response type.

These limitations underscore the inherent challenges of isolating memory-specific neural activity from domain-general processes. While our results argue against a simple executive functional account of the similarity between encoding and retrieval, future work using optimized comparison tasks may further disambiguate the contributions of task-general and memory-specific mechanisms.

Our findings suggest that separate factors underlie the neural features shared between successful encoding and retrieval of verbal memories and the features shared between retrieval and engaged arithmetic problem solving. The key methodology of our study was employing a third comparison task that is relatively dissimilar to both encoding and retrieval. To our knowledge, this approach has not been applied to elucidate the activity common to encoding and retrieval. While cognitive neuroscientists frequently study the relation between the brain and behavior within individual experimental tasks, a fuller understanding of the neural features associated with behavior may be obtained by contrasting activity from multiple tasks engaging a broad array of cognitive functions within the same individuals. Future studies may further leverage cross-task designs with multivariate decoding to better understand subject-level variation in memory systems and its relation to other cognitive processes.

## Methods

### Participants

Patients with pharmacoresistant epilepsy (N=371) were implanted with intracranial electrodes to identify seizure foci as part of routine medical treatment for epilepsy. The presented data were collected as part of a large multi-center study at Columbia University Medical Center (New York, NY), Dartmouth Medical Center (Hanover, NH), Emory University Hospital (Atlanta, Georgia), Hospital of the University of Pennsylvania (Philadelphia, PA), the Mayo Clinic (Rochester, MN), the National Institutes of Health (Bethesda, MD), the University of Colorado Anschutz Medical Campus (Aurora, CO), the University of Texas Southwestern Medical Center (Dallas, TX), and Thomas Jefferson University Hospital (Philadelphia, PA). Each hospital’s institutional review board approved the research protocol, and each participant (or their legal guardian) provided informed consent before data collection began. All participants were screened with standard neuropsychological evaluations for normal cognitive function. Participants were not compensated for their participation.

Prior publications using subsets of our data have presented the electrophysiological correlates of successful memory encoding and retrieval (Burke et al., 2014, 2015; Long et al., 2014) with Kragel et al. (2017) showing significant cross-task decoding between encoding and retrieval in a subset of the data presented here. We note that while our analyses of within-subject cross-task decoding between the math and memory phases of our task are novel, a previous pre-print from our group analyzed between-subject univariate effects of success during the math distractor task along with a statistical model of math problem difficulty (Randazzo et al., Submitted). We provide the de-identified data and code for our analyses at https://memory.psych.upenn.edu/Data.

### Pre-registration of Cross-task Decoding Analyses

To reduce statistical bias from selecting over multiple analysis approaches, we randomly split our full dataset into an exploration dataset, in which we first tuned our methods for cross-task decoding, and a confirmation dataset, in which we subsequently replicated our optimized analyses. We pre-registered our final cross-task decoding analyses with the Open Science Foundation prior to assessing them in the confirmation data (available at https://osf.io/9mg46). Our initial sample consisted of 397 subjects who together completed 1218 sessions of our free recall experiments. Applying our exclusion criteria (see below) resulted in 195 subjects and 574 sessions in the encoding phase, 165 subjects and 437 sessions in the retrieval phase, and 181 subjects and 513 sessions in the math phase in our exploration dataset. For the confirmation set, 175 subjects and 557 sessions were included for encoding, 151 and 420 for retrieval, and 161 and 482 for math respectively. The above numbers include subjects with fewer than three experimental sessions. For our primary cross-task decoding analyses, we exclude subjects with fewer than three experimental sessions each having observations within all three task phases after applying experiment-specific exclusion criteria. This criterion resulted in a sample of 75 subjects and 285 sessions with 63024 encoding events, 13480 retrieval events, and 14515 math events in our exploration dataset, and in our confirmation dataset, 61 subjects and 238 sessions with 55872 encoding events, 11738 retrieval events, and 11974 math events. We present results from the exploration, confirmation, and combined datasets.

### Behavioral Tasks

Participants completed a delayed free recall task in which lists of words were presented on a computer screen one at a time (see Figure 1). The words or study items consisted of common English nouns, with the word pools available at https://memory.psych.upenn.edu/Word_Pools. Subjects completed one or both of two variants of the free recall task: free recall of unrelated words and free recall of categorized words. We pool over these experiment variants in this work. Each list contained 12 items. Study items appeared for 1.6 seconds separated by inter-stimulus intervals uniformly distributed from 0.75 to 1 seconds. Following the study phase, subjects completed a mathematical distractor task, after which they recalled aloud as many of the studied items as they could in any order. The recall phase lasted 30 seconds, and trained annotators manually marked vocalization onsets from experimental audio recordings.

In the mathematical distractor task phase, subjects answered problems of the form “A + B + C = ?” in which A, B, and C were numbers ranging from 1 to 9. Subjects typed their answers and pressed “Enter” to submit their responses. Subjects then received immediate auditory feedback with a high or low-pitched beep indicating whether a response was correct or incorrect respectively. Subjects were sequentially presented problems in this format for 20 seconds. Problems were untimed, and subjects had to complete the final problem in each distractor period even after 20 seconds had elapsed. In each experimental session, patients completed up to 12 or 25 such lists, depending on the variant of the experiment. In some sessions, patients became fatigued and stopped testing before reaching the maximum number of lists per session. Subjects contributed varying numbers of sessions depending on their interest in the experiments and availability to test. In this study, participants included in the cross-task decoding analyses contributed a minimum of three sessions, each with events in all three tasks. In our univariate anatomical analyses, we include participants who completed any number of sessions.

### Statistical Model of Arithmetic Problem Di**ffi**culty

To account for the effect of problem difficulty on subject responses during the arithmetic distractor phase, we modeled log-transformed problem response times using a linear mixed-effects model fit across subjects. We selected model factors based on broad findings in the literature of mathematical cognition (Ashcraft, 1992), on a behavioral model presented in an earlier pre-print from our group using a subset of the data presented here (Randazzo et al., Submitted) (227 subjects reported), and on patterns identified in our exploration data using a forward stepwise selection approach. Our behavioral model included regressors for the problem sum, whether the problem solution was odd, whether any two digits summed to ten, whether any two digits were the same, whether all three digits were the same, and whether a problem required one or two carry operations in addition to random subject-level intercepts and slopes for the problem sum factor and random session-level intercepts. This model was fit to all sessions that passed our exclusion criteria without event-level exclusions for electrophysiological considerations. In particular, we did not enforce the minimum problem RT criterion to avoid strong censorship in the model fit though we retained the maximum problem RT cut-off to exclude outliers. Subject data were included in the model fit regardless of how many sessions a subject completed.

### Electrophysiological Recordings

Patients were implanted subdurally with grid or strip electrodes placed on the surface of the brain, stereotactic depth electrodes placed deep within the brain, or both. Implant locations were selected by physicians solely based on medical considerations. Neighboring recording contacts were typically spaced 10 mm apart for grid or surface electrodes or 5 to 10 mm apart for depth electrodes. Intracranial EEG (iEEG) was recorded using several recording systems depending on the hospital site including clinical monitoring systems (Nihon Kohden EEG-1200, Grass Aura-LTM64, or Natus XLTek EMU 128) and dedicated research systems (Blackrock NSP and the External Neural Stimulator developed by Medtronic). Recordings were either re-referenced to bipolar montages (with channel recordings being subtracted from neighboring channels) or recorded in a bipolar reference mode. We filtered the bipolar re-referenced EEG for line noise using a fourth-order Butterworth notch filter with cut-off frequencies of 58 to 62 Hz. We additionally excluded free recall lists in which the EEG for any channel on any of the list event epochs of a given task phase was constant or otherwise resulted in invalid spectral power values (e.g., from taking the logarithm of zero power).

A small subset of patients were re-implanted or were recorded with multiple recording montages during a hospital stay such that different sets of electrodes were recorded at different times in the same subject. For these analyses, we compute neural decoding scores separately for each montage before averaging scores across montages within subject to obtain the subject-level decoding score. Different montages from a given subject did not cross the exploration-confirmation split.

### Anatomical Localization

Intracranial electrodes were localized using methods described in prior work (Sakon et al., 2022; Herz et al., 2022; Ezzyat et al., 2024). Briefly, our pipeline registered post-implant CT images to pre-implant T1-weighted MRI images using Advanced Normalization Tools (ANTS; Avants et al., 2008). This registration generated coordinates for each electrode in image space which were subsequently transformed to individual-space FreeSurfer coordinates (Fischl et al., 2004). Electrode region labels were assigned using several methods depending on the available information. Clinical neuroradiologists manually annotated regions for a subset of subjects and validated the output of the automated pipeline used to assign region labels to the remaining subjects. The automated pipeline used ANTS and Automated Segmentation of Hippocampal Subfields (ASHS; Yushkevich et al., 2015) to segment whole brain and MTL volumetric regions from the T1-weighted images and hippocampal coronal T2-weighted MRI scans. For electrodes near cortical surface regions for which labels provided by either neuroradiologists or volumetric segmentations were unavailable, surface labels were generated automatically using the Desikan-Killiany-Tourville cortical parcellation protocol from individual-space FreeSurfer coordinates (Fischl et al., 2004; Desikan et al., 2006; Klein and Tourville, 2012). These coordinates were corrected for implant-related brain shift using an energy minimization algorithm in a subset of patients (Dykstra et al., 2012). We localized a bipolar re-referenced pair of recording contacts to the midpoint of the contact coordinates. Finally we excluded somatosensory, motor, and speech-related regions to remove activity related to motor responses during the math distractor and retrieval phases. These regions included pars opercularis, pars triangularis, bank of the superior temporal sulcus, cerebellum, paracentral lobule, supplementary motor cortex, and superior temporal, transverse temporal, precentral, postcentral, and supramarginal gyri.

### Task Decoding

#### Decoding Arithmetic Performance

We decoded subject performance on the arithmetic distractor phase to model engagement in a task involving executive control processes, with no requirement for subjects to encode new memories. Subjects completed the arithmetic task with high accuracy, answering 95.4% of problems correctly (median across subjects in our exploration dataset with an inter-quartile range of 91.9% to 97.6%). To avoid ceiling effects and to create a more balanced classification task than would be obtained from decoding problem correctness, we elected to instead decode whether subjects answered problems faster or slower than their individual median performance within a session. Incorrect trials, trials containing “Backspace” keystrokes, and trials with response times greater than 20 seconds were excluded from our analyses.

While this decoding approach captured subject engagement with the arithmetic task, it does not account for the effects of problem difficulty on response times. We therefore incorporated problem difficulty in a variant of our decoding paradigm using a regression model fit to predict subject response times from characteristics of the arithmetic problems. Using this model, we computed response residuals as the difference between a subject’s response time and the response time predicted by the model for that problem presentation. To account for problem difficulty, we then decoded whether a response residual exceeded or fell below the median response residual for that subject within a given session.

#### Comparing Successful Retrievals to Failed Memory Search

To examine the neural activity associated with episodic retrieval, we contrast successful retrievals against matched periods of silence or “deliberations” indicating moments of failed retrieval during the recall phase (see Figure 1). We define correct retrieval periods as 500 ms epochs preceding correct retrievals subject to the following exclusion criteria. We excluded repeated retrievals and intrusions (vocalizations of words not present in the most recently studied list). Given the importance of the current episodic context to retrieval processes, we limited our study to retrieval events cued by prior retrievals (Howard and Kahana, 2002). We thus excluded the first correct retrieval in each recall phase if it was not proceeded by an intrusion, ensuring each correct retrieval had a well-defined contextual cue. To reduce cross-contamination between the activity of nearby retrievals or other vocalizations, we excluded retrievals preceded by less than 2000 ms of silence. Similarly, we exclude all retrievals having onsets during the first second of the retrieval phase to avoid artifact from the audible beep and fixation cross indicating the start of retrieval.

We define candidate deliberations as periods of silence during the recall phase that meet the following constraints. Candidate deliberation periods must fall after the first recall or intrusion and before the last recall or intrusion in a list. The onset of the preceding vocalization must have occurred at least 2000 ms prior to the onset of the candidate deliberation period, and the onset of the subsequent vocalization must have fallen at least 1000 ms after the onset of the candidate deliberation period. Subject to the above criteria, we selected moments of deliberation within the candidate periods for analysis as follows. For each correct retrieval meeting our inclusion criteria, we attempted to match the vocalization onset time relative to the start of the retrieval phase to a time within a candidate period from a different list. These matches were made exactly when possible and within 5 seconds of the vocalization onset otherwise. When a retrieval could be matched to multiple candidate deliberation periods across different lists (either multiple exact matches if available or else multiple tolerated matches), we selected the match belonging to the list closest to the retrieval, breaking ties randomly. We further required consecutive deliberations to be separated by at least 1000 ms. Retrievals that could not be matched were dropped from subsequent analysis, and we excluded all matches from experimental sessions with fewer than eight matches. These criteria were selected to ensure the onset times and list numbers of retrievals and matched deliberations closely aligned. After applying these criteria, subjects had a median of 25.2% of retrievals matched (inter-quartile range (IQR): 14.8%, 36.9%) in our confirmation data (exploration: 26.7% IQR: 16.4%, 37.0%). Onset times differed between retrievals and deliberations by a median of -7.3 ms (IQR: -76 ms, 258 ms) across subject-level average differences in our confirmation data (exploration data : IQR: -58 ms, 287 ms). List numbers differed by a median of 0.15 (IQR: -0.26, 1.2; exploration: 0.02, IQR: -0.47, 1.1).

### Spectral Feature Extraction

For each task phase, we compute spectral power during event epochs designed to capture moments of successful and unsuccessful cognitive processing (Figure 1A). For the math phase, we computed spectral powers from 1800 ms to 500 ms before the first response keystroke entered by the subject. Given the math problems were untimed, we chose this epoch to coincide with engaged problem solving near the moment of problem resolution. We selected the offset time of this epoch to reduce motor-related artifact generated at the time of response while the epoch duration was selected to maximize predictive performance in our exploration dataset. A mirrored buffer of 500 ms was appended to each end of the math response epochs to further avoid contamination by motor-related artifact. To exclude visual activity related to the onset of problem presentation (which would tend to differentially impact problems with faster responses), we required all EEG epochs to begin at least 500 ms after problem onset.

For the retrieval phase, we analyzed epochs ranging from 600 to 100 ms before the onset of retrieval vocalization or matched deliberation. We used mirror buffers of 500 ms on each side of retrieval epochs and matched deliberation epochs. Mirroring retrieval epochs on the side nearest the vocalization removes vocalization artifact while mirroring both sides of the deliberation intervals prevents overlap between the buffers of nearby deliberations. Finally, to ensure comparisons between retrieval and deliberation epochs are not driven by differences in artifact due to mirror buffering, we similarly mirror buffered the retrieval epochs on the side furthest from the vocalization.

For the encoding phase, we define EEG epochs 1366 ms in length starting from the onset of item presentation as used in prior studies (Ezzyat et al., 2018, 2024). We use mirror buffers of 500 ms for the encoding epochs in line with the arithmetic and retrieval phases.

We computed spectral power across the epochs for each task phase using eight Morlet wavelets with center frequencies logarithmically spaced from 5 Hz to 175 Hz (wave number 5). The resulting wavelet-transformed epoch time series were log-transformed, averaged across time (excluding epoch buffers), and z-scored within experimental session to generate a set of eight spectral features per channel for each epoch.

### Evaluating Neural Decoding of Memory and Mathematical Cognition

We fit subject-level logistic regression classifiers to predict cognitive success in each of the three task phases from neural activity transformed into spectral power features. We evaluated within-task classification performance using cross-validated Area Under the Receiver-Operating Curve (AUC). For evaluating neural decoding, we used the Python scikit-learn library implementations of logistic regression (sklearn.linear model.LogisticRegression with the LIBLINEAR solver and balanced class weighting) and AUC (sklearn.metrics.roc auc score) (Pedregosa et al., 2011; Fan et al., 2008). Each subject included in the neural decoding analyses was required to have completed at least three experimental sessions each with a minimum of 15 events separately for each task passing the relevant exclusion criteria. To ensure our classifiers generalized across experimental contexts, we employed a leave-one-session-out cross validation (LOSO-CV) approach. In LOSO-CV, one session out of the K total sessions for a subject is held out from the model fitting process to provide a test set for unbiased model evaluation. The remaining K-1 sessions are used to fit the model parameters, and the resulting model is evaluated on the held-out session. This process repeats with each session being held out in turn, and the average model performance (AUC) across all held-out sessions provides the final measure of neural decoding performance for a given subject and task. To fit the model parameters for a given held-out session, we similarly used a nested inner LOSO-CV process on the remaining K-1 sessions to optimize the L2 regularization penalty weighting over held-out inner fold sessions, with the linear model parameters for each inner cross-validation fold being fit to K-2 sessions. We optimized the regularization penalty over 52 points logarithmically spaced from 10^−6^ to 10^11^. We then refit the classifier parameters on all K-1 sessions with the optimized regularization weighting before evaluation on the held-out session from the outer cross-validation fold. We averaged AUCs from outer-fold sessions for each subject to obtain the within-task decoding performance for that subject and tested the population average decoding performances for each class against random chance performance of an AUC of 0.5 with one-sample *t*-tests across subjects. As described previously, a small number of subjects were recorded with multiple recording montages. For these subjects, decoding scores were computed separately for each recording montage with at least three sessions and then averaged scores across montages to obtain the decoding score for that subject.

To measure the neural similarity between successful cognition across tasks, we compute the degree to which classifiers fit on a source task A transfer to predicting observations from a target task B. For each subject, we trained a single classifier on all sessions available for task A. The L2 penalty for this classifier was selected to maximize the standard leave-one-session-out cross-validated AUC averaged across hold-out sessions rather than the nested cross-validation approach used for within-task evaluation. We evaluate the predictive performance (AUC) of that subject classifier with the neural features and behavioral outcomes from task B pooled across all sessions to obtain our measure of subject-level task-A-to-task-B cross-task neural decoding. Decoding performance was estimated in this way for all pair-wise combinations of our three cognitive decoding tasks. For each combination of decoding tasks (task A predicting B, or B predicting A), we assess significant performance at the population level using one-sample *t*-tests across subjects compared to random chance performance. We further compare the subject-level decoding performance across task combinations (i.e., whether task A predicts task B better than task C predicts B and whether task B predicts A better than C predicts A) using paired *t*-tests. Specifically we compare the neural similarity between math and retrieval to the similarity between math and encoding by testing math-to-encoding cross-task decoding vs. math-to-retrieval. We also assess the similarities of encoding and retrieval to math with the reversed directions of cross-task decoding by testing encoding-to-math vs. retrieval-to-math. For each direction of cross-task decoding, we match either the source task or the target task across the comparison (for example, in the first comparison listed, the math distractor phase provides the source task for the two cross-task decoding tasks being compared). We analogously compare the similarities of math to encoding and of retrieval to encoding by testing encoding-to-retrieval cross-task decoding vs. encoding-to-math and retrieval-to-encoding vs. math-to-encoding, and we compare the similarities of math and encoding to retrieval by testing retrieval-to-encoding vs. retrieval-to-math and encoding-to-retrieval vs. math-to-retrieval.

We separately correct these two sets of tests of cross-task decoding for multiple comparisons using the Benjamini-Yekutieli method of controlling the false discovery rate (FDR) at a rate of *q* < 0.05 (Benjamini and Yekutieli, 2001). Given cross-task decoding between encoding and retrieval was previously reported in (Kragel et al., 2017), we do not include the tests of encoding-retrieval cross-task decoding against random chance in our FDR correction. We repeat these primary cross-task decoding analyses with two sets of controls. First, we adjust our labels of trial-level arithmetic success for problem difficulty using our behavior model of response times as described previously. Second, we exclude contacts localized to brain regions involved in somatosensation, motor movement, or speech (see above). We assess our results with these controls applied together as well as separately for comparison, basing our primary inferences on the combined control condition.

To control for the effect of within-task decoding strength on these comparisons of cross-task decoding, we repeat a subset of these hypothesis tests after normalizing the cross-task scores by dividing them by the within-task scores for the test task. This analysis was not preregistered. We limit this analysis to subjects with a test task AUC of at least 0.55 to stabilize the normalization. To account for the within-task score of the train task, we compare these normalized scores only for pairs of tasks matched by the train task (e.g., transfer from encoding to retrieval compared to transfer from encoding to math). The resulting three paired *t*-tests (normalized transfer from encoding to retrieval compared to normalized transfer from encoding to math, retrieval-math compared to retrieval-encoding, and math-retrieval compared to math-encoding) are FDR-corrected for multiple comparisons.

To compare our measure of cross-task neural decoding against a more typical measure of neural similarity, we further assess the Pearson correlation between subject-level standardized effects of our cognitive contrasts. For each subject and session we compute Hedge’s *g* of the spectral power between successful and unsuccessful events for every bipolar pair and wavelet frequency. We average these effects across sessions within subject-montage and compute the Fisher z-transformed correlation across the resulting vectors of contact-frequency-level effects for all pairs of tasks. As before, we average these transformed correlations across subject-montages within subject to obtain the final subject-level measure of neural similarity. We test these transformed correlations for each pair of tasks against zero using a one-sample *t*-test. We similarly test the difference in z-transformed correlations between pairs of tasks (e.g., comparing math correlated with retrieval to math correlated with encoding) using paired *t*-tests. We ran these tests separately with the subjects and sessions from the cross-task decoding analysis as well as using all subjects and sessions with data in all three tasks regardless of whether those subjects completed at least three sessions. We also repeated this analysis controlling for arithmetic problem difficulty and motor-related activity as done for the cross-task neural decoding analysis.

### Measuring regional spectral power predicting behavioral outcomes

We computed the spectral activity associated with engaged cognition during each task across several regions of interest (ROIs) used in prior work: superior, middle, and inferior frontal gyri, hippocampus, parahippocampal gyrus, middle and inferior temporal gyri, inferior and superior parietal lobule, and occipital cortex (Ezzyat et al., 2017, 2018). As discussed above, our controls for motor-related activity entirely excluded superior temporal gyrus as well as pars opercularis and pars triangularis from inferior frontal gyrus. Specifically, we computed Hedge’s *g* for each recording electrode and frequency band by comparing successful and unsuccessful events within experimental session for each task (Hedges, 1981). We average these electrode-level effects first within ROIs and frequencies and then across sessions for a given subject. Statistically reliable effects across ROIs and frequencies were inferred with one-sample t-tests across subjects against a null effect of zero. We corrected the resulting p-values for multiple comparisons across ROIs and frequencies using the Benjamini-Krieger-Yekutieli two-stage linear step-up procedure for false discovery rate (FDR) correction at a false discovery rate of *q* < 0.05 (Benjamini et al., 2006).

### Controlling for rehearsal during the math phase

We test the alternative hypothesis that any observed neural similarity between math and retrieval could be due to rehearsal of studied items during the arithmetic distractor phase. If subjects rehearsing during the arithmetic distractor led to 1. slower math response times, 2. higher recall rates, 3. retrieval-related neural activity appearing during the distractor, one could expect spurious cross-task decoding unrelated to neural similarities underlying mathematical cognition and retrieval processes. If this were the case, we would expect that list-level recall rates and math response times would correlate in the same direction as the effect of cross-task decoding. We would also expect that those subject-level correlations would predict cross-task decoding performance. To test this hypothesis, we correlated mean list-level recall rates and response times within all sessions included in our cross-task decoding analyses with at least 10 lists each having at least three math responses. To ensure no biases arose from differing numbers of math responses across distractor phases, we compute the mean RT over only the first three responses during a list. We averaged these session-level correlations within subject after Fisher z-transformation and tested the mean population z-transformed Pearson correlation across subjects against zero. Our exploration data showed no evidence for this rehearsal hypothesis using standard frequentist *t*-tests. To allow for finding evidence in favor of the null hypothesis of no effect of rehearsal, we opted to test this difference in means using a Bayes factor with an objective Jeffreys-Zellner-Siow (JZS) prior for a Cauchy scale factor of 0.5 (Rouder et al., 2009). We further tested the correlation of subject-level recall-RT correlations with cross-task decoding AUCs for math and retrieval against zero. We similarly test this hypothesis using a Bayes factor for correlation with an objective JZS prior (Wetzels and Wagenmakers, 2012). For consistency, we also report the results of the equivalent frequentist *t*-tests for both of these comparisons but base our inferences on the Bayes factors. Finally, these analyses were conducted both with standard RTs as well as with the residualized RTs from our model of problem difficulty.

### Statistical Analyses

All frequentist tests were evaluated at the *p* < 0.05 threshold unless specified otherwise. We corrected for multiple comparisons with false discovery rate (FDR) correction (*p_FDR_* < 0.05; Benjamini and Yekutieli, 2001). Bayes factors were interpreted using standard thresholds, with a Bayes factor of great than 10 (or less than 1/10) providing strong evidence for the alternative (or the null) hypothesis and a Bayes factor between 3 and 10 (or between 1/3 and 1/10) indicating notable but weaker evidence (Wetzels and Wagenmakers, 2012).

## References

1. Arsalidou M, Taylor MJ (2011) Is 2+2=4? Meta-analyses of brain areas needed for numbers and calculations. NeuroImage 54:2382–2393.

2. Ashcraft MH (1992) Cognitive arithmetic: A review of data and theory. Cognition 44:75–106.

3. Avants BB, Epstein CL, Grossman M, Gee JC (2008) Symmetric diffeomorphic image registration with cross-correlation: evaluating automated labeling of elderly and neurodegenerative brain. Medical Image Analysis 12:26–41.

4. Badre D, Wagner A (2007) Left ventrolateral prefrontal cortex and the cognitive control of memory. Neuropsychologia 45:2883–2901.

5. Barredo J, Öztekin I, Badre D (2015) Ventral Fronto-Temporal Pathway Supporting Cognitive Control of Episodic Memory Retrieval. Cerebral Cortex 25:1004–1019 Publisher: Oxford University Press (OUP).

6. Benjamini Y, Krieger AM, Yekutieli D (2006) Adaptive linear step-up procedures that control the false discovery rate. Biometrika 93:491–507.

7. Benjamini Y, Yekutieli D (2001) The control of the false discovery rate in multiple testing under dependency. Annals of statistics pp. 1165–1188.

8. Burke JF, Merkow M, Jacobs J, Kahana MJ, Zaghloul K (2015) Brain computer interface to enhance episodic memory in human participants. Frontiers in Human Neuroscience 8:1055.

9. Burke JF, Sharan AD, Sperling MR, Ramayya AG, Evans JJ, Healey MK, Beck EN, Davis KA, Lucas TH, Kahana MJ (2014) Theta and high–frequency activity mark spontaneous recall of episodic memories. Journal of Neuroscience 34:11355–11365.

10. Cabeza R, Locantore JK, Anderson ND (2003) Lateralization of prefrontal activity during episodic memory retrieval: evidence for the production-monitoring hypothesis. J Cogn Neurosci 15:249–259.

11. Cragg L, Gilmore C (2014) Skills underlying mathematics: The role of executive function in the development of mathematics proficiency. Trends in Neuroscience and Education 3:63–68.

12. Daitch AL, Foster BL, Schrouff J, Rangarajan V, Kasikci I, Gattas S, Parvizi J (2016) Mapping human temporal and parietal neuronal population activity and functional coupling during mathematical cognition. Proceedings of the National Academy of Sciences pp. E7277–E7286.

13. Daitch AL, Parvizi J (2018) Spatial and temporal heterogeneity of neural responses in human posteromedial cortex. Proceedings of the National Academy of Sciences 115:4785–4790.

14. Desikan R, Segonne B, Fischl B, Quinn B, Dickerson B, Blacker D, Buckner RL, Dale A, Maguire A, Hyman B, Albert M, Killiany N (2006) An automated labeling system for subdividing the human cerebral cortex on MRI scans into gyral based regions of interest. NeuroImage 31:968–80.

15. Dykstra AR, Chan AM, Quinn BT, Zepeda R, Keller CJ, Cormier J, Madsen JR, Eskandar EN, Cash SS (2012) Individualized localization and cortical surface-based registration of intracranial electrodes. NeuroImage 59:3563–3570.

16. Ezzyat Y, Wanda P, Levy DF, Kadel A, Aka A, Pedisich I, Sperling MR, Sharan AD, Lega BC, Burks A, Gross R, Inman CS, Jobst BC, Gorenstein M, Davis KA, A. WG, Kucewicz MT, Stein JM, Gorniak RJ, Das SR, Rizzuto DS, Kahana MJ (2018) Closed-loop stimulation of temporal cortex rescues functional networks and improves memory. Nature Communications 9:365.

17. Ezzyat Y, Kragel JE, Burke JF, Levy DF, Lyalenko A, Wanda PA, O’Sullivan L, Hurley KB, Busygin S, Pedisich I, Sperling MR, Worrell GA, Kucewicz MT, Davis KA, Lucas TH, Inman CS, Lega BC, Jobst BC, Sheth SA, Zaghloul K, Jutras MJ, Stein JM, Das SR, Gorniak R, Rizzuto DS, Kahana MJ (2017) Direct brain stimulation modulates encoding states and memory performance in humans. Current Biology 27:1251–1258.

18. Ezzyat Y, Kragel JE, Solomon EA, Lega BC, Aronson JP, Jobst BC, Gross RE, Sperling MR, Worrell GA, Sheth SA, Wanda PA, Rizzuto DS, Kahana MJ (2024) Functional and anatomical connectivity predict brain stimulation’s mnemonic effects. Cerebral Cortex 34:bhad427.

19. Fan RE, Chang KW, Hsieh CJ, Wang XR, Lin CJ (2008) Liblinear: A library for large linear classification. The Journal of Machine Learning Research 9:1871–1874.

20. Fischl B, van der Kouwe A, Destrieux C, Halgren E, Segonne F, Salat DH, Busa E, Seidman LJ, Goldstein J, Kennedy D, et al. (2004) Automatically parcellating the human cerebral cortex. Cerebral Cortex 14:11–22.

21. Gordon AM, Rissman J, Kiani R, Wagner AD (2014) Cortical reinstatement mediates the relationship between content-specific encoding activity and subsequent recollection decisions. Cerebral Cortex 24:3350–3364.

22. Grabner RH, Ansari D, Koschutnig K, Reishofer G, Ebner F, Neuper C (2009) To retrieve or to calculate? Left angular gyrus mediates the retrieval of arithmetic facts during problem solving. Neuropsychologia 47:604–608.

23. Harvey BM, Vansteensel MJ, Ferrier CH, Zuiderbaan W, Aarnoutse E, Bleichner MG, Dijkerman HC, van Zandvoort MJE, Leijten F, Ramsey N, Dumoulin SO (2013) Frequency specific spatial interactions in human electrocorticography: V1 alpha oscillations reflect surround suppression. NeuroImage pp. 424–432.

24. Hedges LV (1981) Distribution theory for glass’s estimator of effect size and related estimators. Journal of Educational Statistics 6:107–128.

25. Herz N, Bar-Haim Y, Tavor I, Tik N, Sharon H, Holmes EA, Censor N (2022) Neuromodulation of visual cortex reduces the intensity of intrusive memories. Cerebral Cortex 32:408–417.

26. Howard MW, Kahana MJ (2002) A distributed representation of temporal contex 47t. Journal of Mathematical Psychology 46:269–299.

27. Jacobs J, Kahana MJ (2009) Neural representations of individual stimuli in humans revealed by gamma-band electrocorticographic activity. Journal of Neuroscience 29:10203–10214.

28. Kahana MJ (2012) Foundations of Human Memory Oxford University Press, New York, NY.

29. Kim H (2011) Neural activity that predicts subsequent memory and forgetting: a meta-analysis of 74 fMRI studies. NeuroImage 54:2446–2461.

30. Klein A, Tourville J (2012) 101 labeled brain images and a consistent human cortical labeling protocol. Frontiers in Neuroscience 6.

31. Knops A, Thirion B, Hubbard EM, Michel V, Dehaene S (2009) Recruitment of an Area Involved in Eye Movements During Mental Arithmetic. Science 324:1583–1585.

32. Kragel JE, Ezzyat Y, Sperling MR, Gorniak R, Worrell GA, Berry BM, Inman C, Lin JJ, Davis KA, Das SR, Stein JM, Jobst BC, Zaghloul KA, Sheth SA, Rizzuto DS, Kahana MJ (2017) Similar patterns of neural activity predict memory function during encoding and retrieval. NeuroImage 155:60–71.

33. Kriegeskorte N, Mur M, Bandettini P (2008) Representational similarity analysis – connecting the branches of systems neuroscience. Frontiers in Systems Neuroscience 2:1 – 28.

34. Long NM, Burke JF, Kahana MJ (2014) Subsequent memory effect in intracranial and scalp EEG. NeuroImage 84:488–494.

35. Lucchelli F, De Renzi E (1993) Primary dyscalculia after a medial frontal lesion of the left hemisphere. *Journal of Neurology*, Neurosurgery & Psychiatry 56:304–307.

36. Manning JR, Polyn SM, Baltuch G, Litt B, Kahana MJ (2011) Oscillatory patterns in temporal lobe reveal context reinstatement during memory search. *Proceedings of the National Academy of Sciences*, USA 108:12893–12897.

37. Marklund P, Fransson P, Cabeza R, Petersson KM, Ingvar M, Nyberg L (2007) Sustained and Transient Neural Modulations in Prefrontal Cortex Related to Declarative Long-Term Memory, Working Memory, and Attention. Cortex 43:22–37 Publisher: Elsevier BV.

38. Pedregosa F, Varoquaux G, Gramfort A, Michel V, Thirion B, Grisel O, Blondel M, Prettenhofer P, Weiss R, Dubourg V, Vanderplas J, Passos A, Cournapeau D, Brucher M, Perrot M, Duchesnay E (2011) Scikit-learn: Machine learning in Python. Journal of Machine Learning Research 12:2825–2830.

39. Pinheiro-Chagas P, Sava-Segal C, Akkol S, Daitch A, Parvizi J (2024) Spatiotemporal dynamics of successive activations across the human brain during simple arithmetic processing. The Journal of Neuroscience p. e2118222024.

40. Polyn SM, Natu VS, Cohen JD, Norman KA (2005) Category-specific cortical activity precedes retrieval during memory search. Science 310:1963–1966.

41. Randazzo M, Ezzyat Y, Kahana MJ (Submitted) Spectral tilt underlies mathematical problem solving. Submitted .

42. Rouder JN, Speckman PL, Sun D, Morey RD, Iverson G (2009) Bayesian t tests for accepting and rejecting the null hypothesis. Psychonomic Bulletin & Review 16:225–237.

43. Sakon JJ, Halpern DJ, Schonhaut DR, Kahana MJ (2022) Hippocampal ripples signal encoding of episodic memories In *Cognitive Neuroscience Society*.

44. Sokolowski HM, Hawes Z, Ansari D (2023) The neural correlates of retrieval and procedural strategies in mental arithmetic: A functional neuroimaging meta-analysis. Human Brain Mapping 44:229–244.

45. Tompary A, Duncan K, Davachi L (2016) High-resolution investigation of memory-specific reinstatement in the hippocampus and perirhinal cortex. Hippocampus 26:995–1007.

46. Tulving E (1979) Relation between encoding specificity and levels of processing In Cermak L, Craik F, editors, Levels of Processing in Human Memory, chapter 19, pp. 405–428. Lawrence Erlbaum Associates.

47. Urgolites ZJ, Wixted JT, Goldinger SD, Papesh MH, Treiman DM, Squire LR, Steinmetz PN (2020) Spiking activity in the human hippocampus prior to encoding predicts subsequent memory. Proceedings of the National Academy of Sciences 117:13767–13770.

48. Vatansever D, Smallwood J, Jefferies E (2021) Varying demands for cognitive control reveals shared neural processes supporting semantic and episodic memory retrieval. Nature Communications 12 Publisher: Springer Science and Business Media LLC.

49. Wagner AD, Schacter DL, Rotte M, Koutstaal W, Maril A, Dale AM, Rosen BR, Buckner RL (1998) Building memories: remembering and forgetting of verbal experiences as predicted by brain activity. Science 281:1188–1191.

50. Weidemann CT, Kragel JE, Lega BC, Worrell GA, Sperling MR, Sharan AD, Jobst BC, Khadjevand F, Davis KA, Wanda PA, Kadel A, Rizzuto DS, Kahana MJ (2019) Neural activity reveals interactions between episodic and semantic memory systems during retrieval. Journal of Experimental Psychology: General 148:1–12.

51. Wetzels R, Wagenmakers EJ (2012) A default Bayesian hypothesis test for correlations and partial correlations. Psychonomic Bulletin & Review 19:1057–1064.

52. Yushkevich PA, Pluta JB, Wang H, Xie L, Ding SL, Gertje EC, Mancuso L, Kliot D, Das SR, Wolk DA (2015) Automated volumetry and regional thickness analysis of hippocampal subfields and medial temporal cortical structures in mild cognitive impairment. Human Brain Mapping 36:258–287.

