## Supplemental Material for "Neural components underlying successful free recall are specific to episodic memory"

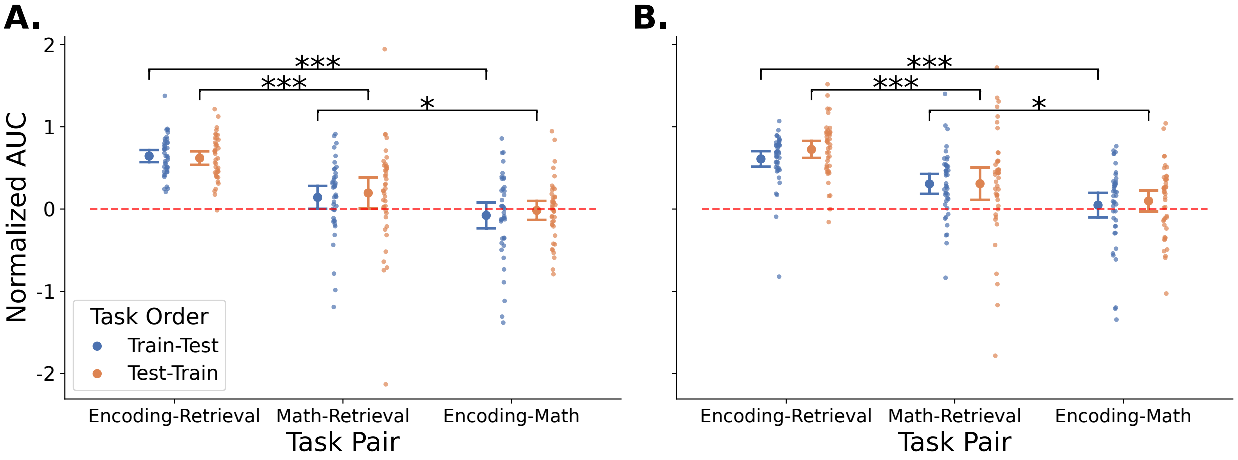


Figure 1: **Distinct neural factors hold after controlling for within-task decoding performance.** We re-evaluate our comparisons of cross-task decoding performance after normalizing by the within-task scores for the test task and only comparing task pairs matched on the train task (see main text). This analysis was not preregistered. A., B. Normalized cross-task decoding between item encoding, spontaneous retrieval, and math distractor performance (A: confirmation data, B: exploration data. Dots show subject-level statistics. Error bars show standard 95% confidence intervals around the mean. Significance bars reflect FDR-corrected paired t-tests. ***: $p < 0.001$, **: $p < 0.01$, *: $p < 0.05$, n.s.: $p \geq 0.05$.

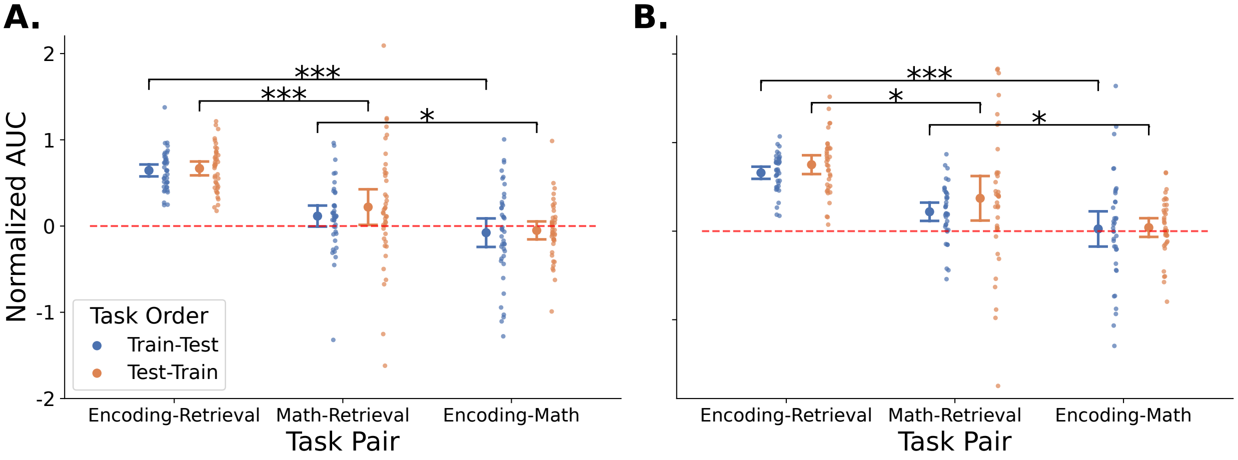


Figure 2: **Distinct neural factors remain after controlling for within-task decoding and math difficulty.** We repeat our cross-task decoding performance analyses now accounting for both within-task decoding performance and arithmetic problem difficulty (see main text). This analysis was not preregistered. Format follows Supplementary Figure 1.
